# RNA and protein are not interchangeable in single-cell spatial multi-omics

**DOI:** 10.64898/2026.10.05.756660

**Authors:** Sergi Cervilla, Daniela Grases, Noelia Montesdeoca, Albert Font, Maria Fernández-Saorin, Olatz Etxaniz, Juan Carlos Pardo, Maria Ochoa-de-Olza, Mariona Figols, Nuria Sala-González, Claudia Fina, Maria Rodriguez, Juan P de Torres, Maria D Lozano, Luis M Montuenga, Carlos de Andrea, Rubén Pio, Vicenç Ruiz de Porras, Eduard Porta-Pardo

## Abstract

Spatial transcriptomics is increasingly used to infer cellular phenotypes, molecular interactions, and functional tissue organization from RNA, although most of these biological processes are ultimately mediated by proteins. Here, we directly tested the interchangeability of spatial RNA and protein measurements using paired Xenium profiling of 21 matched targets across ∼1.8 million cells from primary lung adenocarcinoma and metastatic castration-resistant prostate cancer. After accounting for marker- and dataset-specific protein background, we quantified concordance at multiple levels, including single-cell detection, expression magnitude, cell-type distribution, patient-level abundance, and spatial organization. Agreement at single-cell level was strongly marker-dependent. Canonical lineage markers generally showed concordant RNA and protein patterns, whereas several molecules, including in our study PD-1, PD-L1, and LAG-3, showed little concordance across modalities. Importantly, spatial organization remained substantially more discordant between modalities than overall abundance, indicating that spatial biomarkers are not directly transferable across modalities. Together, our results show that spatial RNA and protein measurements provide largely overlapping information for lineage markers and overall abundance, but can diverse for specific molecules and for higher-order spatial features, where the two layers provide complementary rather than equivalent readouts. These findings provide practical guidance on when spatial transcriptomic readouts can be interpreted as proxies for protein and when orthogonal protein validation is advisable.

## Introduction

Spatially resolved molecular profiling is increasingly used to characterize cellular states, interactions, and tissue organization directly within intact tissues. These studies typically measure one or more molecular layers, most commonly RNA and protein, to describe the biological properties of individual cells and their spatial context. Although transcript and protein abundance are linked through gene expression, their correspondence is imperfect because of differences in translation, degradation, trafficking, localization, and other regulatory processes^1,2^. This relationship has been examined extensively in bulk tissues^3,4^ and, more recently, in dissociated single-cell systems^5,6^. However, incomplete correspondence does not necessarily imply that one modality is an inaccurate representation of the other: RNA and protein capture different molecular aspects of cellular state, and a cell may therefore be positive in one modality and negative in the other. The relevant question is whether these differences are large enough to change the biological conclusions drawn from a tissue.

This becomes particularly important once molecular measurements are interpreted spatially. Spatial analyses increasingly move beyond measuring abundance to ask which cells express a marker, which cell populations are associated with a given state, how expressing cells are organized in tissue, and where putative cellular or molecular interactions occur. In intact tissue, moreover, a detected molecule must not only be measured but also correctly attributed to a segmented cell. Unlike in bulk or dissociated single-cell assays, individual transcripts may be assigned to neighboring cells, membrane-associated protein signal may cross computational cell boundaries, and extracellular or pericellular proteins may not conform to those boundaries at all. Thus, spatial RNA–protein discordance can arise both from genuine differences between molecular layers and from technical differences in how signals are detected and assigned in situ.

Despite these additional complexities, RNA–protein agreement in spatial data has generally been evaluated using relatively simple measures of correspondence. Several spatial multi-omic studies have reported variable relationships between RNA and protein measurements^7,8^, but these observations have largely emerged from technology demonstrations, small datasets, or analyses focused on overall expression correlation. Whether the two modalities provide equivalent answers across the different levels at which spatial data are interpreted remains less clear. Agreement in overall tissue abundance, for example, could coexist with disagreement about which individual cells are positive, which cell types carry the signal, or how that signal is spatially organized. This distinction may be especially consequential for molecules such as immune checkpoints, for which transcript and protein measurements are both biologically meaningful but describe different molecular states and may therefore support different interpretations of the same tissue.

Here, we directly evaluated RNA-protein concordance using simultaneous Xenium profiling of matched RNA and protein targets in two independent cancer cohorts comprising lung adenocarcinoma and metastatic castration-resistant prostate cancer. Across approximately 1.8 million cells, we assessed agreement at multiple levels, including binary detection, expression magnitude, cell-type ranking, patient-level abundance, and spatial organization. We find that concordance is strongly marker-dependent, with agreement lower at the single-cell and spatial levels than for overall abundance. These results show that RNA and protein provide overlapping and complementary but non-interchangeable representations of tissue biology, and that the molecular layer measured should be matched to the biological question being asked.

## Results

### Paired spatial RNA and protein profiling across two cancer cohorts

To directly compare RNA and protein measurements within the same spatial context, we generated simultaneous RNA – protein profiles from two independent cancer cohorts (**Figure 1a**). We used the Xenium Immuno-Oncology panel (targeting 380 RNA molecules), along with a 27-plex protein panel. There are 21 markers shared in both modalities (**Figure 1b**). Within this overlap, the protein panel resolves CD45RA and CD45RO isoforms, whereas the transcript panel captures only the shared parent gene, *PTPRC*. Similarly, the CD16 protein corresponds to *FCGR3A* at the transcript level. Therefore, we consider both cases as expected sources of RNA–protein discordance rather than technical artifacts from now on.

**Figure 1:**
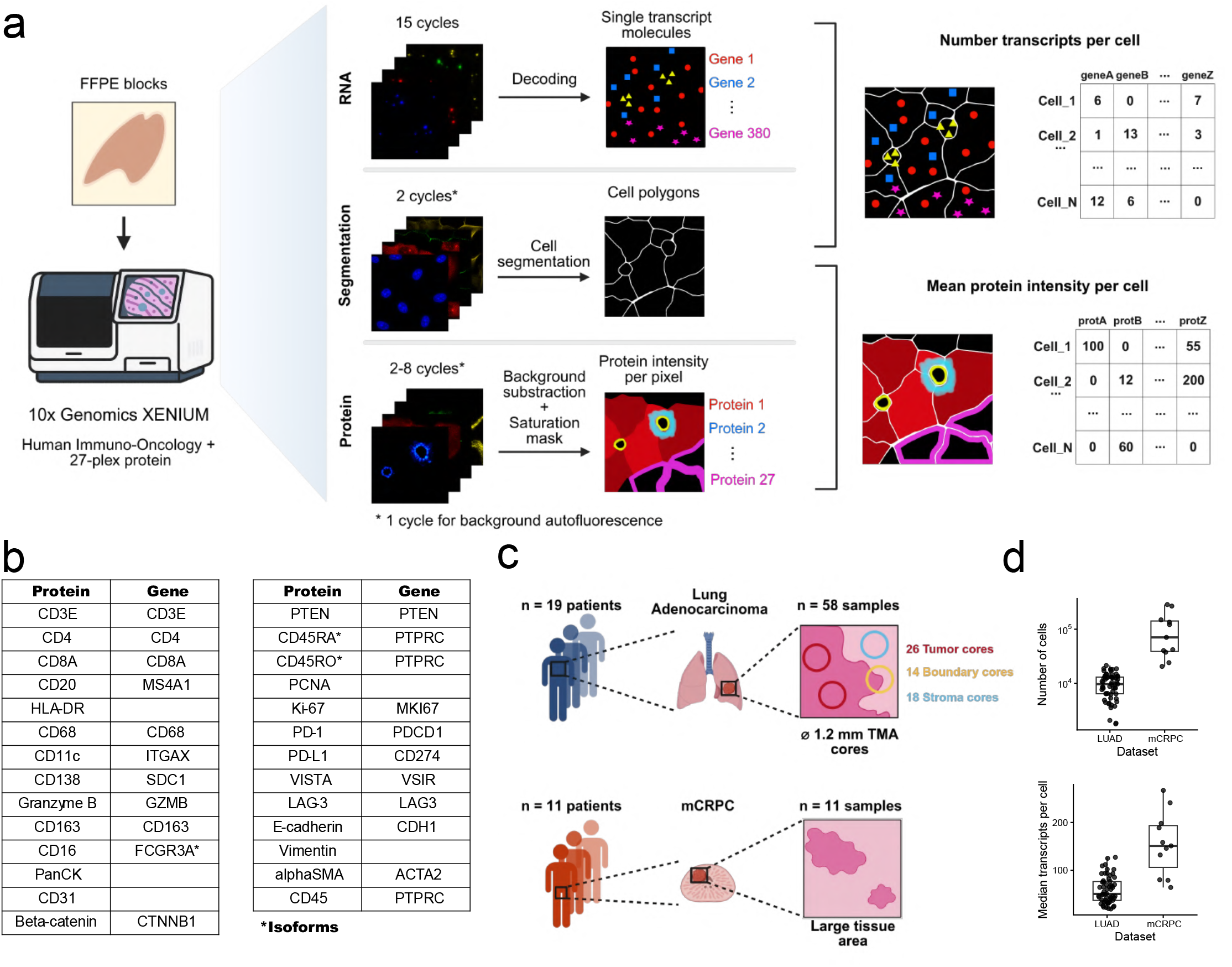
Xenium multi-omics platform overview and dataset summary. **(a)** Overview of the Xenium multi-omics data acquisition workflow. **(b)** Correspondence between the protein sub-panel and the Xenium Immuno-Oncology gene panel; empty cells indicate protein markers with no matching gene target. **(c)** Summary of the generated multi-omics datasets. **(d)** Boxplot showing number of cells per sample (top) and median number of transcripts per cell (bottom) in each dataset.

The first cohort comprised primary tumors from 19 treatment-naive patients with lung adenocarcinoma (LUAD), sampled as tissue microarray (TMA) cores of 1.2 mm diameter, with up to four cores per patient spanning tumor, boundary, and stroma regions, yielding 58 cores that passed quality control (**Figure 1c**). The second cohort includes single large tissue sections from 11 patients with metastatic castration-resistant prostate cancer (mCRPC) who had received prior taxane-based chemotherapy (**Figure 1c**). The LUAD dataset was generated in 2025 as part of the Xenium multi-omics early-access program, whereas the mCRPC cohort was generated independently in-house in 2026. The study was conducted in accordance with the Declaration of Helsinki and the protocols approved by the respective institutional ethics committees (2019.111, and PI-21-353).

After filtering low-quality cells and artifact regions (**Methods**), we retained 1.77M cells in total, with a median of 9,654 cells per sample in LUAD and 70,555 per sample in mCRPC, reflecting the smaller tissue area sampled per LUAD core relative to the single large mCRPC section (**Figure 1d**). Median transcripts per cell were 50.5 in LUAD and 151 in mCRPC, within the range reported in previous Xenium benchmarking studies using targeted gene panels^9–11^ (**Figure 1e, Figure S1, Table S1**). Together, these datasets provide two independent tissue contexts in which matched RNA and protein measurements can be compared from individual cells to higher-order spatial organization.

### Establishing interpretable protein measurements for cross-modal comparison

Before assessing RNA-protein concordance, we first examined the behavior of protein measurements independently. The reason is that, whereas the RNA readouts from Xenium experiments have been extensively benchmarked, the protein measurements are relatively new, and potentially subject to issues such as background, spillover or marker-specific sensitivity that could otherwise be mistaken for biological discordance.

For most markers, such as CD8A, the mean per-cell intensity distribution is zero-inflated, with a large fraction of cells showing no detectable signal, and the remaining non-zero values broadly distributed across the intensity range without a distinct second mode (**Figure 2a**). This lack of a clearly separated positive and negative population precluded the use of simple mixture-based thresholds to define protein-positive cells. Additionally, non-zero intensity did not necessarily indicate true expression, as it could be caused by other factors, such as the membrane-associated signal extending across segmentation boundaries and contributing intensity to neighboring cells (**Figure 2b**).

**Figure 2:**
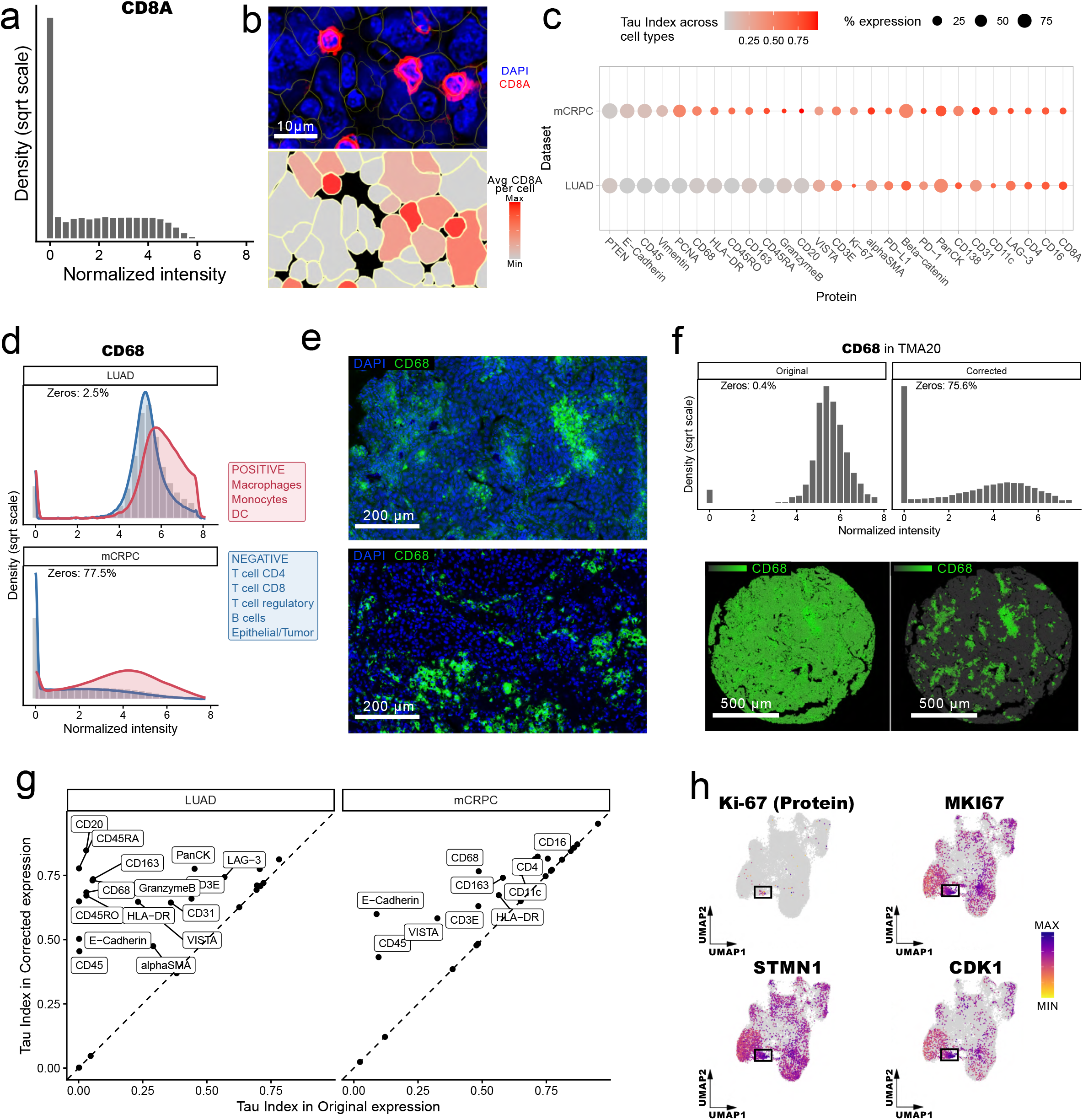
Protein detection quality, specificity, and background correction. **(a)** CD8A protein intensity distribution. Histogram of normalized mean CD8A protein intensity per cell. **(b)** CD8A spatial expression. Top: CD8A protein staining (red) and DAPI nuclear staining (blue), with cell-segmentation boundaries overlaid in yellow, in an enlarged region of a LUAD sample. Bottom: spatial distribution of the normalized mean CD8A protein intensity per cell. **(c)** Cell-type specificity of protein markers. Dot plot showing protein expression specificity quantified using the Tau index across 18 cell types, in the LUAD (n = 58 samples) and PROST (n = 11 samples) datasets. Color indicates the Tau index, and point size indicates the percentage of cells with non-zero protein expression. **(d)** CD68 intensity distribution by cell-type identity. Histograms of normalized mean CD68 protein intensity per cell, with overlaid densities (square-root transformed) of expected positive populations in red (Macrophages, Monocytes and DC) and negative populations (T cell CD4/CD8/Reg, B cells and Epithelial/Tumor) in each dataset. **(e)** CD68 staining across datasets. CD68 protein staining (green) and DAPI nuclear staining (blue), displayed using the full slide-level intensity range, in LUAD (top) and PROST (bottom). **(f)** Effect of background correction on CD68. Top: histogram of normalized mean CD68 protein intensity per cell in a LUAD core using raw intensities (left) and background-corrected intensities (right). Bottom: corresponding spatial distributions of CD68 protein expression in the LUAD tissue core. **(g)** Specificity before and after background correction. Comparison of protein-expression specificity (Tau index) calculated using raw mean intensities (x-axis) and background-corrected mean intensities (y-axis) in each dataset. **(h)** Proliferation marker concordance at single-cell resolution. UMAP embedding of cells from a single patient with LUAD, colored by Ki-67 protein abundance and the RNA expression of MKI67, STMN1, and CDK1.

To evaluate how well protein intensity reflects expected cell-type identity, we cross-referenced non-zero expression against cell types inferred independently from the RNA modality (**Methods**) and quantified marker specificity using Tau index (**Figure 2c**). Some differences in Tau specificity between datasets are expected, since LUAD and mCRPC represent distinct tumor biologies that may differ in the cell type composition and their cell-type-specific expression pattern of a given marker, independent of any technical variation.

Most markers showed expression restricted to their expected cell types. However, Vimentin, PCNA, CD68, HLA-DR, CD163, CD45RA, Granzyme B, and CD20 all showed lower Tau specificity in LUAD than in mCRPC, coinciding with a high fraction of non-zero-expressing cells (>90%) for these markers in LUAD. For lineage-restricted markers such as CD68, CD163, CD20, and Granzyme B, this widespread low-level signal was inconsistent with their expected biological distribution and suggested dataset-specific protein background, suggesting dataset-specific protein background, possibly reflecting pre-analytical factors.

As an example, the CD68 intensity distribution differs markedly between datasets. In mCRPC, it follows a zero-inflated distribution, in which the signal is preferentially coming from macrophages, monocytes, and dendritic cells. In LUAD, by contrast, low but non-zero, CD68 signal extended across many cell types that included both, expected positive and negative populations (**Figure 2d, Figure S2**). Inspection at the pixel level confirmed widespread low-intensity fluorescence throughout LUAD tissue, whereas high-intensity signal remained concentrated in putative CD68-positive cellular aggregates (**Figure 2e**). This pattern was not observed in mCRPC, where background and positive signal were more clearly separated.

To reduce this dataset- and marker-specific background, we estimated an empirical baseline for each protein. Briefly, we used cell populations expected to lack that marker, and subtracted this value from the raw per-cell intensity, setting negative values to zero (**Methods**). After this correction, the normalized CD68 intensity distribution in LUAD converged toward the pattern observed for CD68 and other specific lineage markers in mCRPC, and the spatial distribution of signal became consistent with expected marker localization (**Figure 2f**). Applying this correction across both datasets, for all markers with an identifiable negative population, increased Tau specificity overall, with the largest gains in the LUAD markers listed previously as problematic (**Figure 2g**).

Another problematic marker in LUAD was Ki-67/*MKI67*, as only 0.2% of LUAD cells showed non-zero protein intensity, compared to >10% in mCRPC. To determine whether this low detection reflects technical dropout or a genuinely low proliferative fraction in LUAD, we compared protein intensity against gene expression profiles. Using unsupervised clustering of gene expression from the TMAs of a single patient, we identified a cluster of proliferating/cycling cells. This cluster showed high expression of *MKI67* transcript and co-expression of other proliferation markers, including *STMN1* and *CDK1*, despite the corresponding cells showing no detectable Ki-67 protein intensity (**Figure 2h, Table S2**).

Together, these analyses identified two distinct technical features of the protein modality: widespread background signal that could be empirically corrected, and marker-specific loss of sensitivity that could not. We therefore analyzed the RNA–protein comparisons using only background-corrected protein intensities, while interpreting markers with evidence of modality-specific technical failure separately.

### RNA-protein concordance is strongly marker-dependent at the single-cell level

We next asked whether RNA and protein measurements identify the same molecular state in the same cells. Because the two modalities differ fundamentally in how signal is captured (RNA is detected as discrete single-molecule spots that are counted per cell to yield transcript counts, whereas protein is measured as immunofluorescence intensity, averaged across all pixels within a cell) we did not expect their absolute values to be directly comparable (**Figure 3a**). Instead, we evaluated the agreement between both modalities at two levels: detection concordance (whether a cell is called positive or negative for a marker consistently across modalities), and magnitude concordance (whether the quantitative level of signal aligns between modalities once a cell is called positive in both). We defined overall agreement for each gene-protein pair using four metrics: detection concordance (Cohen’s Kappa), magnitude correlation across the whole tissue (Spearman coefficient), magnitude correlation restricted to cells positive in both modalities (Spearman coefficient), and spatial agreement (Lee’s L) (**Figure 3b, Figure S3**).

**Figure 3:**
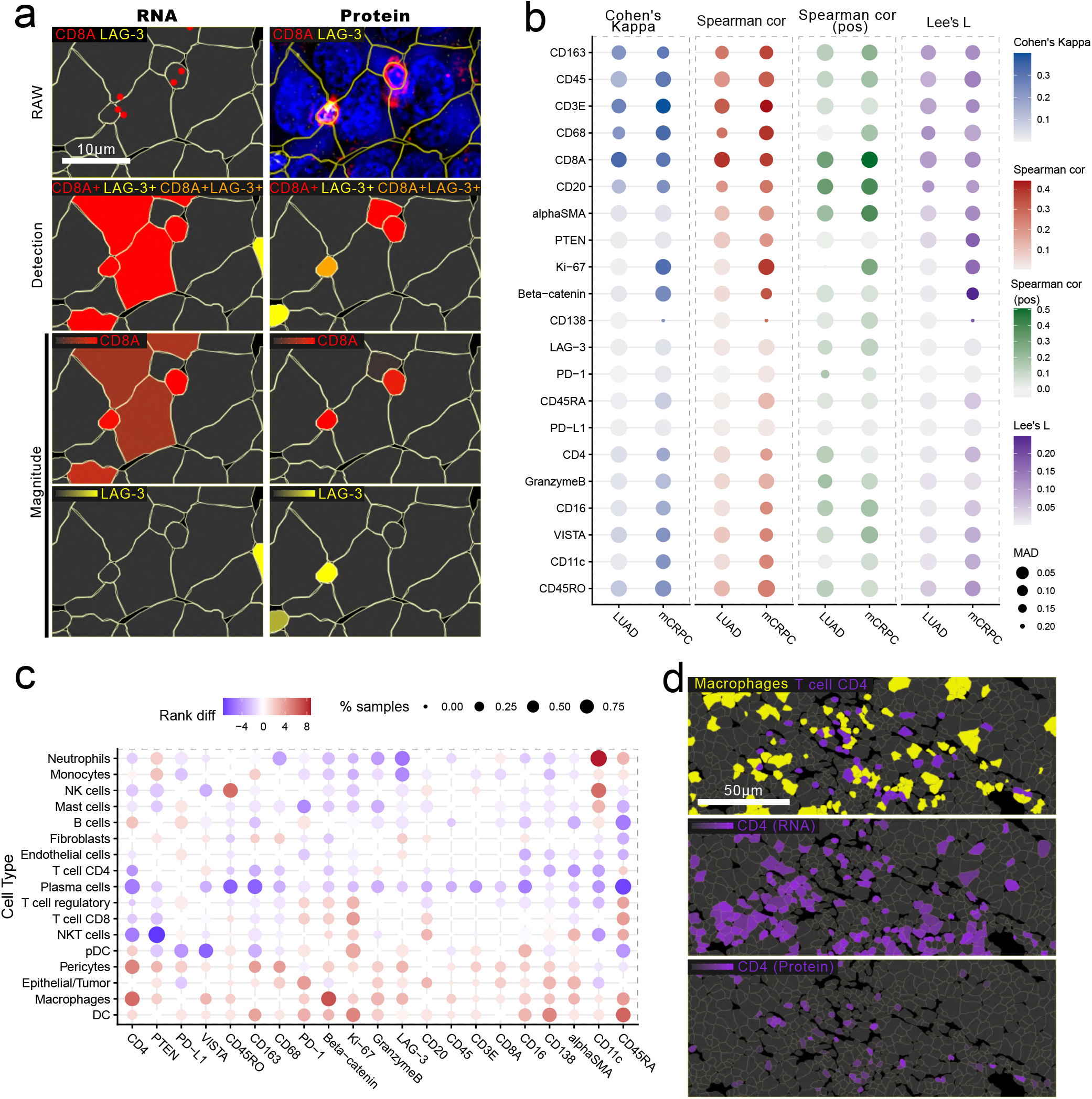
RNA–protein concordance at single-cell level. **(a)** Single-cell RNA and protein detection. Comparison of RNA (left) and protein (right) detection in an enlarged region of a LUAD sample. From top to bottom: (1) raw detection of individual mRNA molecules, shown as colored dots, and protein signal, shown as pixel-level intensity; (2) categorical detection based on non-zero per-cell expression, with cells classified as CD8A+ (red), LAG-3+ (yellow), or CD8A+LAG-3+ (orange); (3) normalized per-cell CD8A expression; (4) normalized per-cell LAG-3 expression. **(b)** Summary of concordance metrics. Dot plot summarizing RNA–protein concordance in each dataset. The evaluated metrics were detection agreement (Cohen’s κ), Spearman’s correlation across all cells, Spearman’s correlation among positive cells, and Lee’s L spatial concordance. Color indicates the median value of each concordance metric, and point size indicates its median absolute deviation (MAD). **(c)** Cell-type ranking discrepancies between modalities. Dot plot showing the ranking difference between modalities for each cell type by mean expression. Dot size indicates the number of samples in which that cell type shows a rank discrepancy of at least 5; color indicates the median rank difference, with red indicating higher rank in RNA and blue indicating higher rank in protein. **(d)** CD4 RNA versus protein specificity. Enlarged region of a LUAD sample, colored by cell type (top; Macrophages in yellow, T cell CD4 in purple) and by normalized per-cell expression of CD4 RNA (middle) and CD4 protein (bottom).

Agreement varied substantially across markers, and the results were broadly reproducible across the two cancer cohorts. Because median transcripts per cell differed ∼3-fold between LUAD and mCRPC, we downsampled mCRPC to match LUAD’s transcript detection level, this did not alter concordance patterns (**Figure S4-S7**), indicating that the cross-cohort differences described below are not driven by differences in per-cell transcript capture. Canonical immune lineage markers (CD163, CD45, CD3, CD68, CD8A, CD20) showed high concordance across all four metrics in both datasets. Within this group, CD8A and CD20 showed comparatively higher magnitude agreement when restricted to double-positive cells. On the other hand, PTEN and alpha-SMA showed no detection concordance, yet their magnitude and spatial agreement remained high, suggesting a mismatch in the threshold used to call positive cells rather than a true absence of agreement in signal intensity. Beta-catenin and Ki-67 showed high agreement only in mCRPC. For Ki-67, this is consistent with the detection failure described above for LUAD. A second group of markers, CD4, Granzyme B, CD16, VISTA, CD11c, and CD45RO, showed lower agreement scores than the lineage markers but remained positive, primarily in mCRPC. In contrast, CD138, CD45RA, PD-1, PD-L1, and LAG-3 showed little or no concordance across the evaluated metrics in either cohort. The discordance of CD45RA was expected in part because the RNA panel measures the shared PTPRC transcript instead of the CD45RA isoform itself.

Finally, we asked whether RNA-protein discordance is influenced by cell-type identity, beyond the marker-specific technical explanations described above, that is, whether certain cell types regulate the relationship between transcript and protein abundance differently. We quantified this using the ranking difference between modalities across cell types and found that macrophages, pericytes, dendritic cells, and epithelial cells were consistently ranked higher by RNA than by protein, indicating that transcript abundance overstates relative expression in these populations compared to their protein counterpart (**Figure 3c**). CD4 in macrophages was among the clearest examples of this pattern, consistent with known post-translation regulation of CD4, including receptor internalization and surface downregulation, in myeloid cells^12^ (**Figure 3d**).

Overall, these results indicate that RNA–protein concordance is not a property of the assay or of the tissue but, instead, it is strongly marker-dependent: for some molecules, the two modalities identify similar cells and spatial patterns, whereas for others they provide substantially different representations of the same tissue.

### Sample-level RNA-protein concordance does not extend to spatial organization

The consistent lack of agreement for PD-1, PD-L1, and LAG-3 is particularly relevant because these markers have directly matched transcript and protein targets and showed discordance independently in both cancer types. In the LUAD TMA, for example, we observed not only differences in abundance within tumor region delineations but also in spatial organization: LAG-3 shifted from a dispersed pattern in RNA to hot-spot clustering in protein (**Figure 4a**).

**Figure 4:**
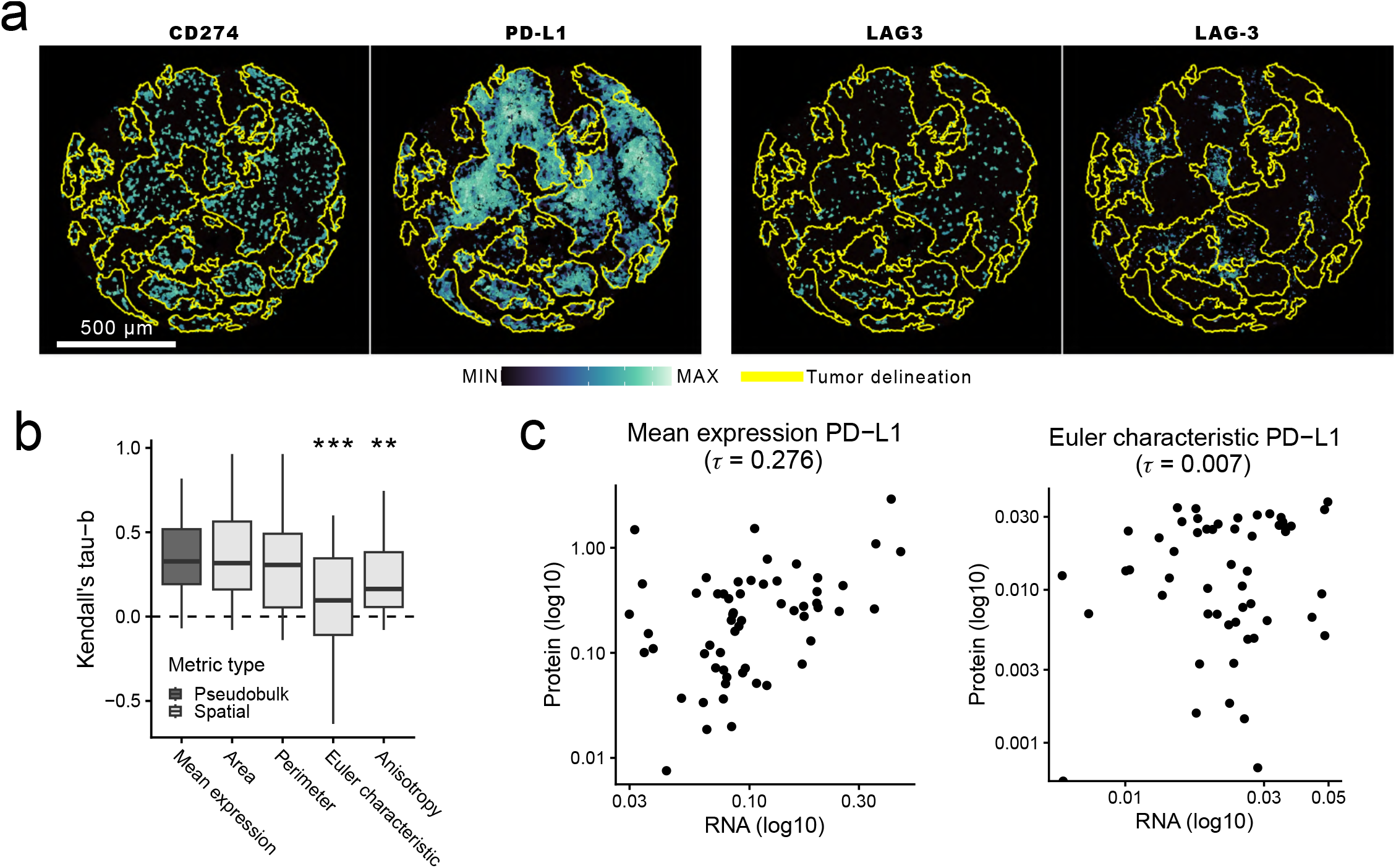
Spatial pattern metrics improve RNA–protein rank concordance for checkpoint markers. **(a)** Spatial expression of checkpoint markers. Spatial expression of CD274 RNA (first), PD-L1 protein (second), LAG3 RNA (third), and LAG-3 protein (fourth) in a LUAD sample; tumor regions are delineated by yellow polygons. **(b)** Concordance across summary metrics. Boxplot of Kendall’s tau-b coefficients quantifying rank concordance between modalities across samples for each metric: mean expression (pseudobulk), area, perimeter, Euler characteristic, and anisotropy (spatial). Each box shows the distribution of tau-b values across markers for that metric. Significance of differences relative to mean expression was assessed with paired Wilcoxon test (**P < 0.01, ***P < 0.001). **(c)** PD-L1 rank concordance by metric. Comparison of sample-level RNA–protein rank concordance for PD-L1 using mean expression (left) and Euler characteristic (right) as the summary statistic per sample, in the LUAD dataset. RNA (x-axis) and protein (y-axis) values are shown on a log10 scale. τ indicates Kendall’s rank correlation between RNA and protein sample rankings.

Given the clinical relevance of these findings for biomarker-based patient stratification, we explored RNA-protein agreement across the full marker panel, comparing ranking concordance based on aggregated abundance, as previously used for both modalities in biomarker studies^13^, against ranking concordance based on spatial features derived from the same data, which we expected to be more sensitive to the single-cell discordances described above.

For spatial organization, we used Minkowski functionals and tensors, simple geometric descriptors widely used in astrophysics and recently applied to spatial gene expression^14^, to quantify area, perimeter, Euler characteristic (a measure of connectivity, e.g., isolated islands versus a networked structure with holes), and anisotropy (directional organization). Mean expression, our bulk-equivalent metric, showed ranking agreement between modalities similar to area and perimeter (Kendall∼0.4), both of which are closely tied to the number of expressing cells (**Figure 4b, Figure S7**). Euler characteristic and anisotropy, in contrast, showed significantly lower agreement between modalities than the bulk-equivalent metric (p = 2.21 × 10^−6^ and p = 9.21 × 10^−4^, respectively). Specifically, patient ranking by PD-L1 mean intensity (bulk-equivalent) was concordant between RNA and protein except for six samples, whereas ranking by Euler characteristic diverged substantially for the same samples (**Figure 4c**).

In conclusion, patient ranking was more consistent between RNA and protein when based on aggregated abundance. Spatial metrics derived from the same data, however, were not interchangeable between modalities, yielding different answers about how the signal is spatially organized within the tissue.

## Discussion

Spatial transcriptomics is increasingly used not only to map gene expression, but to reconstruct the functional organization of tissues from RNA. Yet many of the biological entities being inferred, including cell-surface receptors, ligands, immune checkpoints and signaling molecules, ultimately act as proteins. Here, by measuring RNA and protein in the same cells across two independent cancer cohorts, we show that the spatial information provided by these two molecular layers is complementary, and that their equivalence depends on the marker and on the level of analysis. Concordance varied markedly across markers and according to how agreement was defined: canonical lineage markers frequently showed similar patterns across modalities, whereas other molecules showed little agreement in detection, abundance or spatial organization. Importantly, aggregation could make the modalities appear more concordant even when they disagreed at the individual-cell level or in their higher-order spatial organization. Identifying the contexts in which both layers converge is therefore essential for the robust interpretation of spatial data. Conversely, divergence between modalities should not automatically be interpreted as contradictory: because biological, technical and pre-analytical factors can all contribute to RNA–protein discordance, divergent results may also provide complementary and functionally relevant information on tissue biology.

This problem has characteristics that are specific to spatial in situ measurements. In bulk analyses, RNA and protein are averaged across large populations of cells, potentially masking discordance occurring at the cellular level. Conversely, in dissociated single-cell assays, cells are physically separated before measurement, substantially reducing ambiguity as to which cell a detected molecular signal belongs. In intact tissue, a molecule must not only be detected but also correctly attributed to a segmented cell. Membrane-associated protein signal can extend across segmentation boundaries^15^, individual transcripts can be assigned to neighboring cells^16^, and extracellular or pericellular protein does not necessarily conform to computationally defined cellular territories. Our results illustrate both modality-specific detection problems, including marker-dependent protein background and sensitivity, and local cases in which cellular assignment can influence the resulting readout. Spatial RNA–protein discordance can therefore arise from at least three distinct sources: modality-specific detection, spatial attribution, and genuine molecular biology. This additional attribution problem means that conclusions from bulk or dissociated single-cell RNA–protein comparisons cannot simply be transferred to in situ spatial multi-omics.

The consequences are particularly relevant for immune checkpoints. PD-1, PD-L1 and LAG-3 were among the least concordant RNA–protein pairs in both cohorts, with discrepancies extending from individual-cell detection to spatial organization. *CD274, PDCD1* and *LAG3* transcripts are commonly used in single-cell and spatial transcriptomic studies to define checkpoint-associated cell states, identify putative checkpoint-expressing populations and infer spatial ligand–receptor interactions^17^. Our results do not imply that these transcripts are biologically uninformative; rather, they show that their spatial distribution should not automatically be interpreted as the distribution of the corresponding therapeutic protein. An RNA-negative cell may still be protein-positive, just as RNA positivity does not guarantee detectable protein. This distinction is particularly important for spatial inference: two modalities may provide similar estimates of overall abundance while giving different answers about which cells carry the molecule, whether expressing cells form isolated or connected structures, or where putative molecular interactions occur. Spatial transcriptomics therefore directly measures a transcriptional state; inference of the corresponding protein state remains an additional biological assumption. Notably, variable concordance has already been reported in clinical settings both between PD-L1 immunohistochemistry assays using different antibody clones^18^ and between PD-L1 protein and mRNA in situ hybridization in lung cancer^19^. Our observations should therefore be interpreted in the context of the specific antibody clones included in the Xenium protein panel.

At the same time, discordance should not necessarily be viewed exclusively as measurement error. This is distinct from the cases described above in which discordance could be attributed to a specific technical cause, such as the loss of Ki-67 protein sensitivity in LUAD, consistent with the known sensitivity of Ki-67 immunoreactivity to the age of archival FFPE material^20,21^, or the isoform-level mismatch underlying CD45RA/RO. For markers such as PD-1, PD-L1 and LAG-3, no such explanation was evident from our data and discordance was reproduced in both cohorts. Beyond technical factors, RNA and protein may instead represent different stages of molecular biology operating over different timescales^22^. In some contexts, RNA-positive/protein-negative cells could reflect cells entering a state before the corresponding protein accumulates, whereas RNA-negative/protein-positive cells could reflect persistence of protein after transcription has ceased. Protein can also have a spatial provenance that RNA cannot reveal. Whereas transcripts generally identify the cell in which they were synthesized, proteins may be secreted, shed, deposited extracellularly, or transferred between cells through mechanisms including extracellular vesicles or membrane exchange^23–25^. A protein signal detected within or around a segmented cell therefore need not always have originated from that cell. Our cross-sectional data cannot distinguish these possibilities from residual technical dropout or other forms of post-transcriptional regulation, and direct experimental validation will be required. Nevertheless, for these specific markers, the findings raise the possibility that RNA–protein discordance may sometimes encode cellular history or intercellular communication, rather than representing only noise to be removed.

These findings have practical implications for the choice of spatial profiling technology. Our results do not argue that either transcriptomics or proteomics is universally superior. Whole-transcriptome RNA measurements provide much greater molecular breadth and are therefore particularly powerful for unbiased discovery of cell states, pathways and unexpected biological programs. Protein measurements more directly interrogate the molecular products that execute many cellular functions and may therefore be preferable when the question concerns a defined receptor, ligand, therapeutic target or phenotype. Multimodal profiling is most valuable when the correspondence between these layers is not known: its purpose is not simply to add more features, but to establish which molecular layer is required to answer a particular biological question. Once this relationship has been established, large-scale validation or eventual clinical implementation may be better served by the simplest and least costly assay that directly measures the relevant endpoint.

## Limitations

Our study has several limitations. First, the two cohorts differ in tumor type, anatomical site, year of data generation, sampling strategy, and transcript depth. Therefore, their differences (including the elevated protein background observed in LUAD and the generally higher RNA–protein concordance in mCRPC, to which deeper transcript capture may contribute) cannot be attributed to any single factor and are best interpreted as observations across distinct experimental and biological contexts rather than as technical replicates.

Second, pre-analytical factors may have contributed to the technical behavior of the LUAD dataset. The LUAD tissue microarrays were constructed from archival FFPE blocks obtained between 2008 and 2016, whereas the mCRPC samples were collected more recently, between 2021 and 2025. Prolonged storage of FFPE material progressively reduces RNA integrity and can compromise the antigenicity of certain epitopes, with Ki-67 immunoreactivity being particularly sensitive to the age of archival material^20,21^. Lung tissue also exhibits substantial intrinsic autofluorescence, which may have contributed to the diffuse low-level protein background observed in this cohort.

Third, the performance of individual antibodies within this specific platform may not be optimal for all targets. The antibodies included in the Xenium protein panels are applied in a highly multiplexed workflow whose tissue pretreatment and antigen recognition conditions are shared with RNA detection rather than optimized for each epitope. Limited specificity, suboptimal sensitivity or elevated non-specific binding for some antibodies could therefore contribute to low-level background, reduced detection or apparent RNA-protein discordance. Moreover, the clones used may differ from those validated for clinical immunohistochemistry, which is particularly relevant for PD-L1, for which different clinical assays are known to yield non-equivalent staining^26^.

Fourth, cell-type annotations were derived from the RNA modality and were subsequently used to define expected negative populations for protein-background correction. This approach is more robust for canonical lineage markers with well-established restricted expression, but becomes less certain for markers or cellular states whose expression is not tightly coupled to cell identity. Our concordance estimates are also influenced by analytical choices: binary positivity based on a zero threshold is sensitive to detection efficiency and zero inflation, as is correlation calculated across all cells, making correlations within double-positive cells more directly interpretable to compare expression magnitude.

Finally, our study is restricted to a targeted set of matched markers and two tumor contexts. Most importantly, neither modality provides an independent ground truth. Without orthogonal measurements, cross-sectional data cannot determine whether individual instances of RNA–protein discordance arise from transcriptional or post-transcriptional biology, differences in molecular persistence or localization, or modality-specific detection and attribution errors. Our conclusions are therefore limited to whether RNA and protein provide concordant measurements and biological interpretations, rather than which modality is correct.

## Methods

### Xenium multi-omics sample preparation

The Xenium workflow began by sectioning 5μm FFPE tissue sections onto the Xenium slides, according to the “Xenium FFPE Tissue Preparation Guide” (CG000578 Rev F, 10X Genomics). Briefly, the sections were cut, floated in an sterilized miliq water bath at 42ºC, and carefully placed onto the capture area of a Xenium slide (PN-3000941). After sectioning, the slides with tissue sections were incubated at 42ºC for 3 hours and the slides with sections from TMAs were incubated at 42ºC overnight, after that were kept in a sealed bag with desiccators at room temperature.

Xenium slides with Lung TMA sections were sent to the 10x Genomics laboratory (Pleasanton, CA, USA) for processing, under a Catalyst program and a Software Beta Agreement established between 10x Genomics and the research group at IJC.

Xenium slides with Prostate Cancer Sections were deparaffinized, decrosslinked, and permeabilized prior to probe hybridization following the Xenium In Situ Gene and Protein Expression with Cell Segmentation Staining User Guide (CG000819 Rev A).

Spatial gene expression was profiled using the pre-designed Xenium Human Immuno-Oncology Profiling Panel (v1 chemistry, 380 genes). Genes were selected to enable identification of major immune cell types and subtypes and include immune checkpoint molecules, supporting characterization of the tumor microenvironment at the cellular and subcellular level. Each pre-designed panel additionally includes 40 negative control codewords and 20 negative control probes to assess non-specific signal and decoding accuracy.

Simultaneously, protein detection was performed using the Xenium Protein assay, which is compatible with all Xenium v1 chemistry human gene expression panels. Cell segmentation markers (E-cadherin, Vimentin, AlphaSMA, and CD45) and Xenium Protein subpanels (Immune Cell A/B/C, Immune Checkpoint, Tumor and Proliferation & Differentiation).

Probe hybridization, washing, ligation, and rolling circle amplification for RNA targets were performed with the Xenium In Situ Gene and Protein Expression with Cell Segmentation Staining User Guide (CG000819 Rev A), and the slides were loaded into the Xenium analyzer following User Guide (CG000584 Rev K). The instrument provided a sample scan with nuclei detection, the areas of interest were selected and the analysis began. Following RNA decoding and cell segmentation imaging cycles, seven additional imaging cycles were performed to capture protein subpanel signal. A second background image was acquired for background subtraction on the protein channel prior to quantification.

For each field of view, image data were captured across multiple Z-planes with a 0.75 µm step size spanning the full tissue thickness, across all fluorescence channels per cycle, which were subsequently processed and stitched to generate a spatial map of protein signal.

### Xenium Preprocessing

Gene and protein expression matrices generated by the Xenium Analyzer (v4.0.0.18) were loaded into Seurat (v5) and filtered to retain cells with > 3 transcripts.

For LUAD datasets, cells were assigned to their corresponding TMA core using QuPath’s^27^ “TMA de-array” function (default settings, core size 1.3 mm) applied to the morphology_focus image of each slide. The resulting core polygon coordinates were exported as GeoJSON, imported into R with the sf package^28^, and used to assign each cell centroid to its corresponding TMA polygon. Individual Seurat objects were then created by splitting the dataset by TMA. Cores were excluded if more than 80% of the tissue area was missing or the spatial structure was substantially disrupted; regions containing artifacts or tissue folding were also excluded.

Gene expression was log-normalized using Seurat’s default normalization. Protein expression was normalized using an inverse hyperbolic sine (asinh) transformation with a cofactor of 10.

### Cell annotation

Cell type labels were assigned using a Seurat label transfer approach against a public single-cell reference datasets^29,30^ using the RNA modality. The reference was subset to genes present in the Xenium immuno-oncology panel, and low-quality reference cells were removed by excluding cell types not characterized by the subset of genes and cells requiring a minimum of 10 total RNA counts per cell. The reference’s fine-grained cell type labels were collapsed into coarse cell type categories to match the resolution of expected populations in both datasets; pericytes were assigned separately based on the reference’s finer cell type annotation (**Table S3**). The reference and spatial datasets were then log-normalized, and the top variable features were used to scale the data and compute a PCA embedding.

Label transfer was performed using Seurat’s FindTransferAnchors/TransferData workflow. Anchors were identified between the reference and each spatial query object using the set of genes shared between the reference and the Xenium panel, with the reference PCA (30 dimensions) as the shared embedding. Coarse cell type labels were then transferred to each spatial dataset using TransferData over the same dimensions. Cells with a prediction score lower than 0.4 were assigned as “Unknown”.

### Expression specificity of protein across cell types

For each protein marker, cells were classified as expressing using a > 0 threshold on background-corrected and the percentage of expressing cells was computed per cell type and dataset. Cell-type specificity was summarized using the tau index, where τ approaches 0 for markers expressed broadly across cell types and 1 for markers restricted to a single cell type.

### Background correction

Background thresholds for each protein marker were estimated empirically using cell populations expected to be negative for that marker, defined per-marker in (**Table S4**). For each protein marker, cell types were grouped according to their expected marker status (positive or negative) (**Table S5**), and cell type groups with fewer than 5 cells were excluded. For cell types annotated as marker-negative, the 90th percentile of expression was computed per cell type, and the marker’s background threshold for that sample was defined as the 75th percentile of these per-cell-type 90th percentile values. This procedure was repeated independently for every sample, and the final background threshold for each marker was defined as the median of the per-sample thresholds across the full dataset. Thresholds were subtracted from raw mean intensities and negative values clipped to 0.

### Identification of the proliferative cluster and differential expression

Xenium cells from the TMA cores of a single patient (patient 1) were subset from the merged LUAD dataset. Highly variable genes were selected with Seurat’s FindVariableFeatures (default settings), excluding genes with zero counts, and dimensionality reduction was performed using sparse quasi-Poisson PCA on Pearson residuals (scPearsonPCA), with residuals clipped at 10 standard deviations and genes centered and scaled. The resulting 30 dimensions of the PCA embedding was used for UMAP visualization, construction of a shared nearest-neighbor graph, and Louvain clustering with Seurat’s FindClusters (default resolution). The proliferative cluster (cluster 10) was identified from the cluster-level marker genes (FindAllMarkers). Differential expression of the proliferative cluster against all remaining cells was then assessed with Seurat’s FindMarkers (default Wilcoxon rank-sum test).

### Quantifying RNA–protein concordance

For each sample, RNA and protein signals were compared using transcript counts log-normalization and mean protein intensity were background-corrected an inverse hyperbolic sine (asinh) transformation, cofactor of 10.

RNA–protein concordance was assessed using four complementary metrics per marker pair. First, cells were classified as RNA-positive or protein-positive using a > 0 threshold on normalized values, and detection agreement was quantified with Cohen’s kappa between the two binary calls. Second, expression magnitude concordance was assessed with Spearman correlation, computed both across all cells and restricted to double-positive cells (RNA-positive and protein-positive), to separately capture overall and positive-population-specific concordance. Third, spatial co-association was evaluated using Lee’s L statistic using a spatial neighbor graph (10 nearest neighbors of each cell) and 99 Monte Carlo permutations. Marker pairs with fewer than 3 valid observations were excluded from correlation calculations.

### Transcript depth downsampling

To control differences in transcript depth across datasets, a transcript per cell downsampling was applied in the mCRPC dataset to match LUAD. Raw Xenium transcript counts were then downsampled to a common target median depth of 50 transcripts/cell using scuttle’s binomial thinning (downsampleMatrix), with a distinct random seed per replicate (20 replicates per sample) and re-normalized after thinning. For each depth-matched replicate, the spatial Minkowski profile, global per-marker concordance (detection kappa, Spearman correlation, Lee’s L) metrics described above were recomputed.

### Spatial texture analysis

For each TMA core, cell centroids were binned onto a shared square grid with a bin size of 20 µm, and a binary tissue mask was defined as bins containing at least one cell of any type. For each Protein marker and Xenium gene, mean expression per bin was computed by averaging normalized expression values of cells falling within each bin; bins containing no cells were assigned a value of 0. Expression fields were binarized at a fixed threshold (density ≥ 1) prior to functional computation.

Scalar Minkowski functionals, defined as area fraction (W0), perimeter (W1), and Euler characteristic (W2), were computed from each binarized field using a 2×2 pixel-configuration lookup table, restricted to the tissue mask via minus-sampling to avoid spurious boundary contributions from the mask edge, and normalized by total tissue area.

Local anisotropy was quantified using the Minkowski boundary tensor W_1_^{0,2}, computed as the length-weighted contour integral of the outer product of the local boundary normal with itself, traced along isocontours of the expression field at the binarization level. Eigenvalues of the tensor (λmax, λmin) were used to define the anisotropy index and the dominant eigenvector defined the boundary orientation angle.

### Sample ranking concordance

To assess whether RNA-protein concordance rankings across samples were preserved for spatial texture metrics relative to pseudobulk expression, per-marker Minkowski profiles were aggregated across all samples and reshaped into paired RNA-protein values per metric (mean expression, W0, W1, W2, β) for each marker-gene pair. Rank agreement between assays was quantified using Kendall’s tau-b, computed pairwise-complete across samples within each pair and dataset. To test whether spatial metrics showed reduced rank concordance relative to pseudobulk expression, Kendall’s tau-b values for each spatial metric were compared against mean expression using paired Wilcoxon signed-rank tests across marker pairs.

## Funding

mCRPC sample collection and data generation was funded by the ISCIII grant, project n° PI21/00199. LUAD sample collection and data generation was funded by CIBERONC (CB16/12/00443), the Alberto Palatchi Foundation, and the Instituto de Salud Carlos III, co-funded by the European Union (PI23/00573 and AC24/00153). S.C. was supported by the FI-STEP predoctoral fellowship from the Department of Research and Universities of the Generalitat de Catalunya and co-funded by the European Social Fund Plus (2025 STEP 00466). E.P-P. is supported by supported by the Agencia Estatal de Investigación (AEI, Spain) and the Ministerio de Ciencia, Innovación y Universidades (MICIU, Spain) under grant PID2024-159258OB-I00 / MCIN/AEI/10.13039/501100011033 and Fundació Josep Carreras.

## Notes

### Competing Interest Statement

The authors have declared no competing interest.

