## Supplemental Figures for "RNA and protein are not interchangeable in single-cell spatial multi-omics"

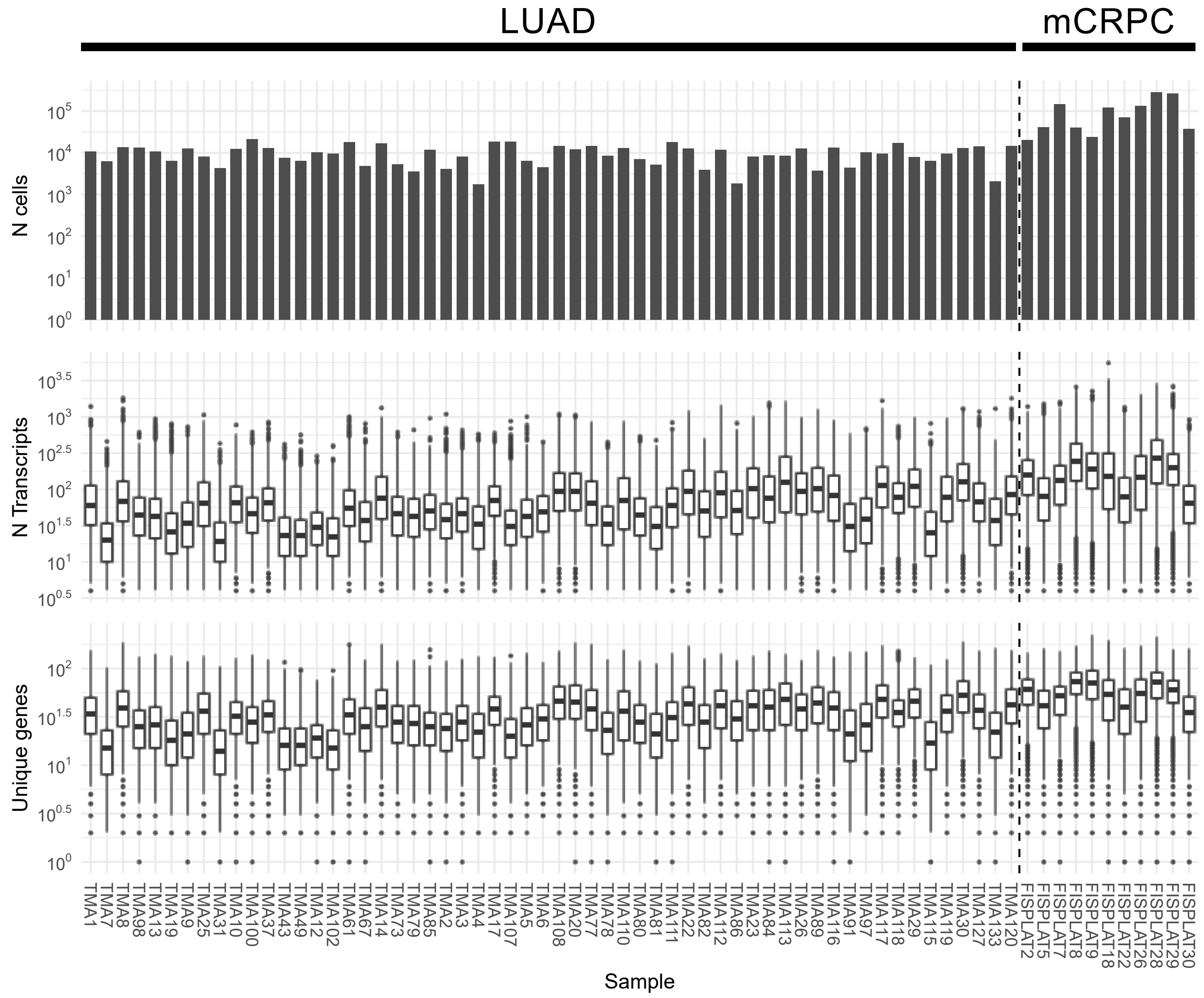


**Figure S1: QC transcriptomics summary.** Summary of the number of cells (top), transcripts per cell (middle) and unique genes per cell (bottom) in each sample in LUAD and mCRPC.

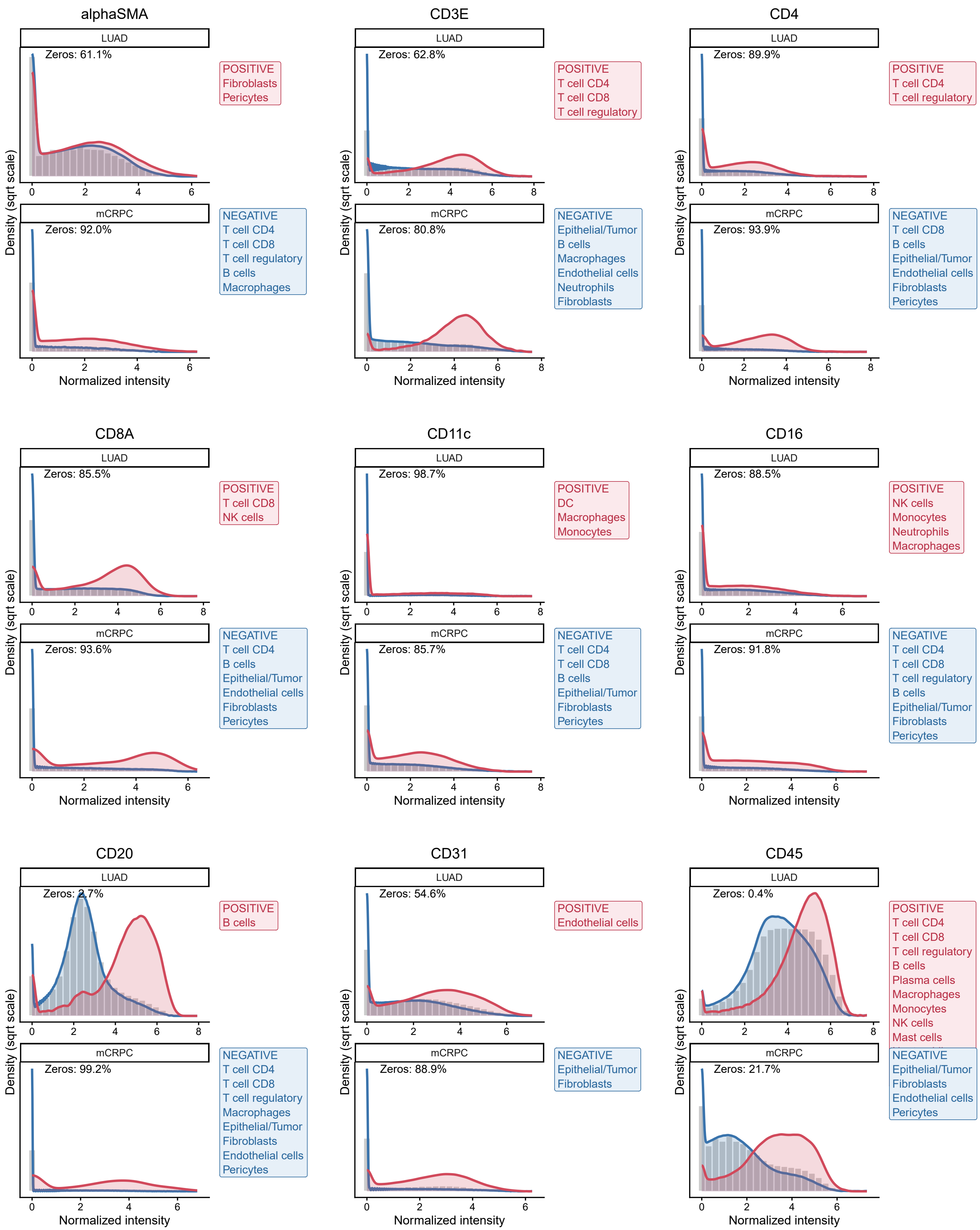

alphaSMA
LUAD
Zeros: 61.1%
POSITIVE
Fibroblasts
Pericytes
mCRPC
Zeros: 92.0%
NEGATIVE
T cell CD4
T cell CD8
T cell regulatory
B cells
Macrophages
CD3E
LUAD
Zeros: 62.8%
POSITIVE
T cell CD4
T cell CD8
T cell regulatory
mCRPC
Zeros: 80.8%
NEGATIVE
Epithelial/Tumor
B cells
Macrophages
Endothelial cells
Neutrophils
Fibroblasts
CD4
LUAD
Zeros: 89.9%
POSITIVE
T cell CD4
T cell regulatory
mCRPC
Zeros: 93.9%
NEGATIVE
T cell CD8
B cells
Epithelial/Tumor
Endothelial cells
Fibroblasts
Pericytes
CD8A
LUAD
Zeros: 85.5%
POSITIVE
T cell CD8
NK cells
mCRPC
Zeros: 93.6%
NEGATIVE
T cell CD4
B cells
Epithelial/Tumor
Endothelial cells
Fibroblasts
Pericytes
CD11c
LUAD
Zeros: 98.7%
POSITIVE
DC
Macrophages
Monocytes
mCRPC
Zeros: 85.7%
NEGATIVE
T cell CD4
T cell CD8
B cells
Epithelial/Tumor
Fibroblasts
Pericytes
CD16
LUAD
Zeros: 88.5%
POSITIVE
NK cells
Monocytes
Neutrophils
Macrophages
mCRPC
Zeros: 91.8%
NEGATIVE
T cell CD4
T cell CD8
T cell regulatory
B cells
Epithelial/Tumor
Fibroblasts
Pericytes
CD20
LUAD
Zeros: 2.7%
POSITIVE
B cells
mCRPC
Zeros: 99.2%
NEGATIVE
T cell CD4
T cell CD8
T cell regulatory
Macrophages
Epithelial/Tumor
Fibroblasts
Endothelial cells
Pericytes
CD31
LUAD
Zeros: 54.6%
POSITIVE
Endothelial cells
mCRPC
Zeros: 88.9%
NEGATIVE
Epithelial/Tumor
Fibroblasts
CD45
LUAD
Zeros: 0.4%
POSITIVE
T cell CD4
T cell CD8
T cell regulatory
B cells
Plasma cells
Macrophages
Monocytes
NK cells
Mast cells
mCRPC
Zeros: 21.7%
NEGATIVE
Epithelial/Tumor
Fibroblasts
Endothelial cells
Pericytes
Density (sqrt scale)
Normalized intensity

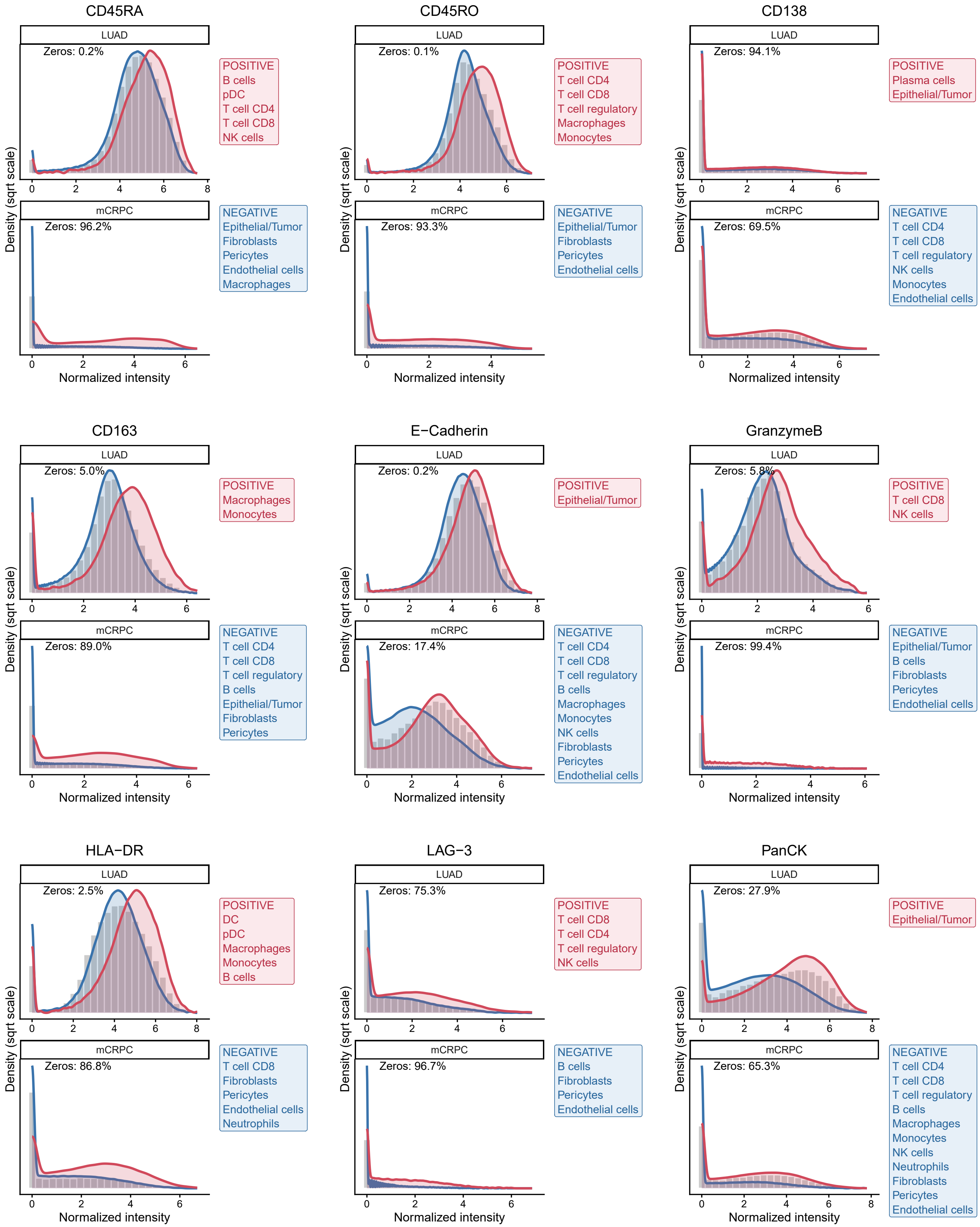
CD45RA
LUAD
Zeros: 0.2%
POSITIVE
B cells
pDC
T cell CD4
T cell CD8
NK cells
mCRPC
Zeros: 96.2%
NEGATIVE
Epithelial/Tumor
Fibroblasts
Pericytes
Endothelial cells
Macrophages
CD45RO
LUAD
Zeros: 0.1%
POSITIVE
T cell CD4
T cell CD8
T cell regulatory
Macrophages
Monocytes
mCRPC
Zeros: 93.3%
NEGATIVE
Epithelial/Tumor
Fibroblasts
Pericytes
Endothelial cells
CD138
LUAD
Zeros: 94.1%
POSITIVE
Plasma cells
Epithelial/Tumor
mCRPC
Zeros: 69.5%
NEGATIVE
T cell CD4
T cell CD8
T cell regulatory
NK cells
Monocytes
Endothelial cells
CD163
LUAD
Zeros: 5.0%
POSITIVE
Macrophages
Monocytes
mCRPC
Zeros: 89.0%
NEGATIVE
T cell CD4
T cell CD8
T cell regulatory
B cells
Epithelial/Tumor
Fibroblasts
Pericytes
E−Cadherin
LUAD
Zeros: 0.2%
POSITIVE
Epithelial/Tumor
mCRPC
Zeros: 17.4%
NEGATIVE
T cell CD4
T cell CD8
T cell regulatory
B cells
Macrophages
Monocytes
NK cells
Fibroblasts
Pericytes
Endothelial cells
GranzymeB
LUAD
Zeros: 5.8%
POSITIVE
T cell CD8
NK cells
mCRPC
Zeros: 99.4%
NEGATIVE
Epithelial/Tumor
B cells
Fibroblasts
Pericytes
Endothelial cells
HLA−DR
LUAD
Zeros: 2.5%
POSITIVE
DC
pDC
Macrophages
Monocytes
B cells
mCRPC
Zeros: 86.8%
NEGATIVE
T cell CD8
Fibroblasts
Pericytes
Endothelial cells
Neutrophils
LAG−3
LUAD
Zeros: 75.3%
POSITIVE
T cell CD8
T cell CD4
T cell regulatory
NK cells
mCRPC
Zeros: 96.7%
NEGATIVE
B cells
Fibroblasts
Pericytes
Endothelial cells
PanCK
LUAD
Zeros: 27.9%
POSITIVE
Epithelial/Tumor
mCRPC
Zeros: 65.3%
NEGATIVE
T cell CD4
T cell CD8
T cell regulatory
B cells
Macrophages
Monocytes
NK cells
Neutrophils
Fibroblasts
Pericytes
Endothelial cells
Density (sqrt scale)
Normalized intensity

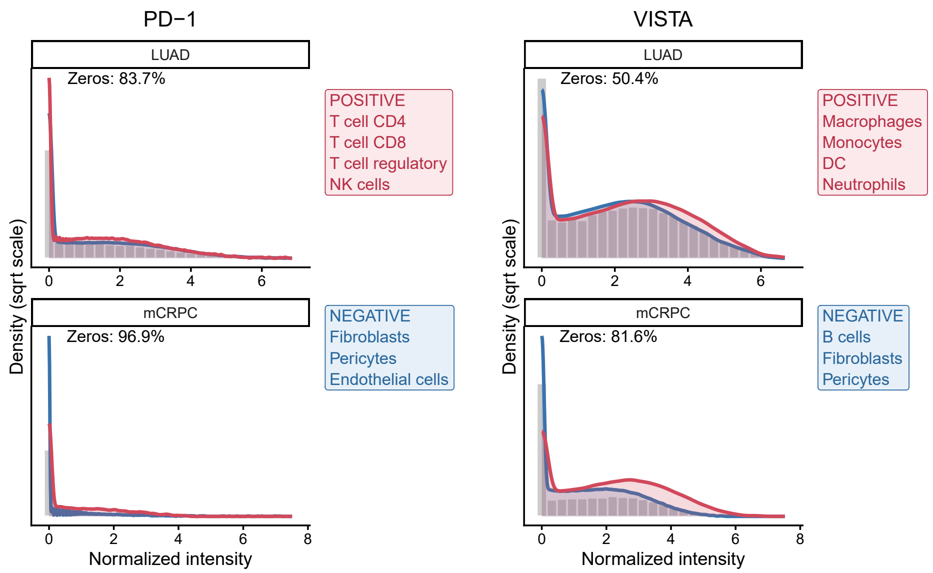


**Figure S2: Mean protein intensity distribution.** Histograms of normalized mean protein intensity per cell (square rooted transformed), with overlaid densities of the expected positive (red) and negative (blue) cell-type populations, for each protein marker in the LUAD and mCRPC datasets.

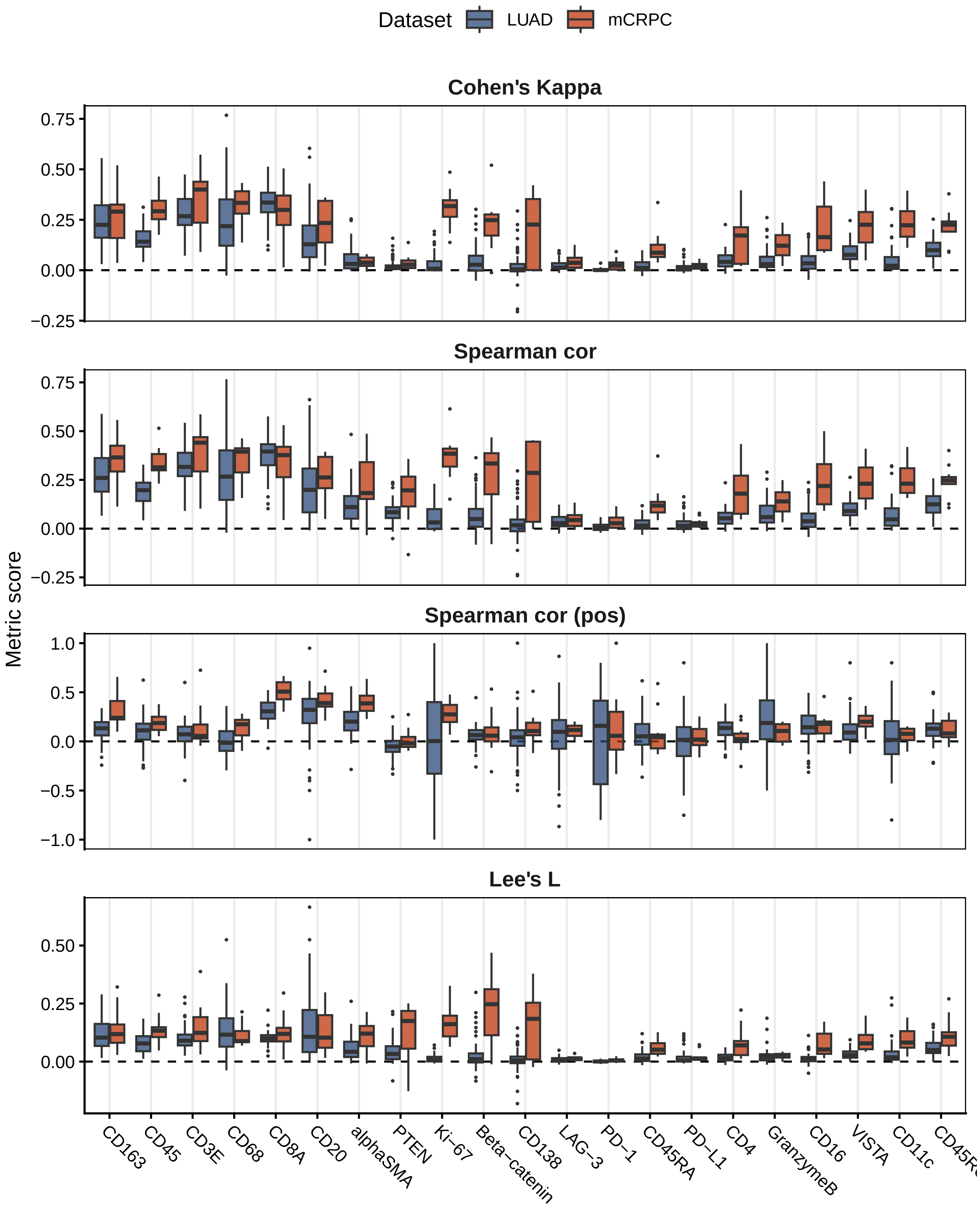


**Figure S3: RNA-protein concordance at single-cell level.** Distribution of RNA-protein concordance metrics in each sample of LUAD (blue) and mCRPC (orange) datasets. The evaluated metrics were detection agreement (Cohen's κ), Spearman's correlation across all cells, Spearman's correlation among positive cells, and Lee's L spatial concordance.

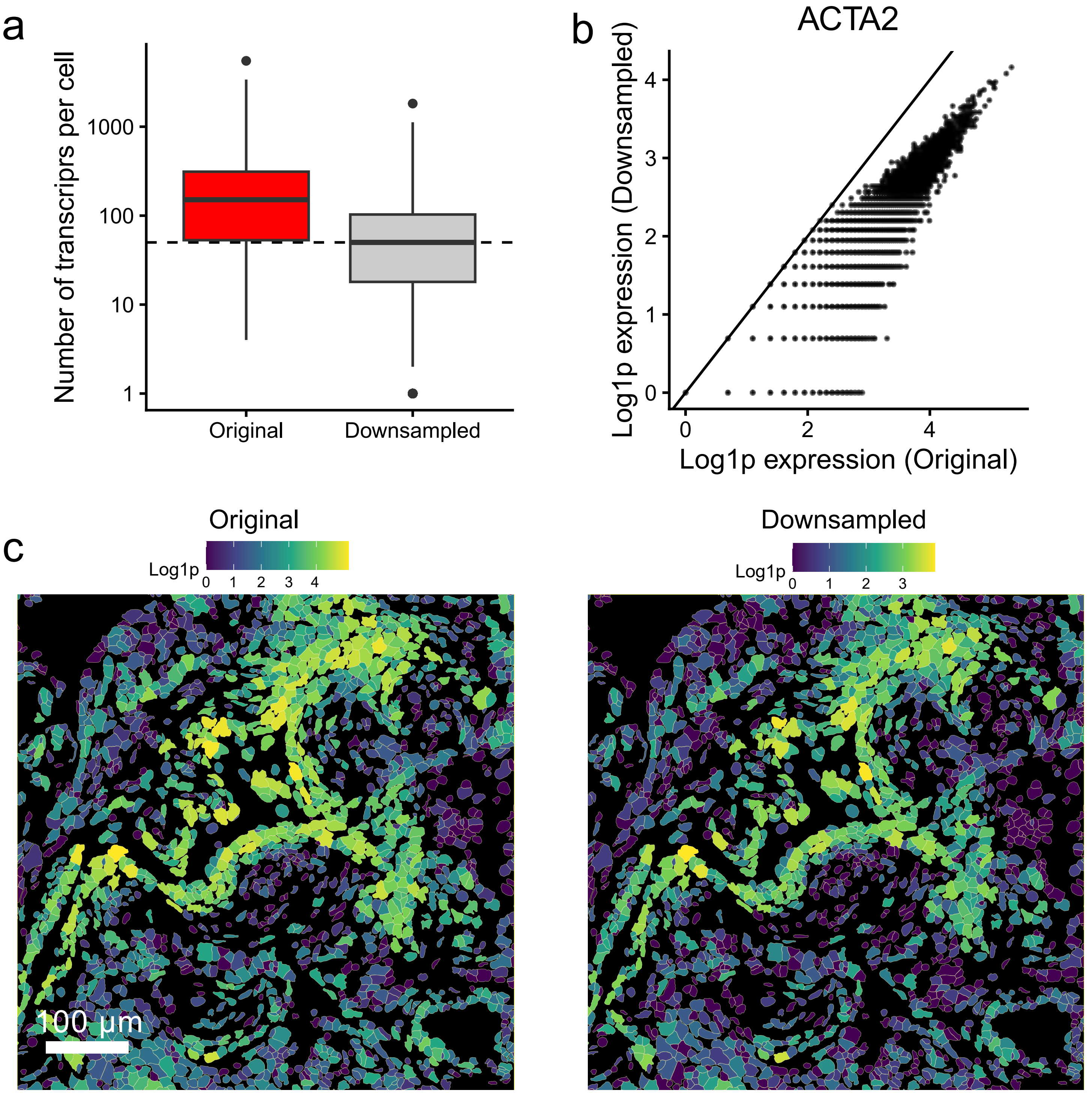


**Figure S4: Assessment of transcript downsampling.** Number of transcripts per cell distributions of FISPLAT18 (mCRPC) at original transcript depth (red) and after downsampling the number of transcripts similar to LUAD median (horizontal dashed line) Log1p ACTA2 expression per cell at original transcript depth (x-axis) versus after downsampling (y-axis) in the FISPLAT18 sample. The diagonal line indicates the expected perfect correlation (x = y). Enlarged region of a FISPLAT18 (mCRPC) sample showing the Log1p ACTA2 expression at original transcript depth (left) and after downsampling (right).

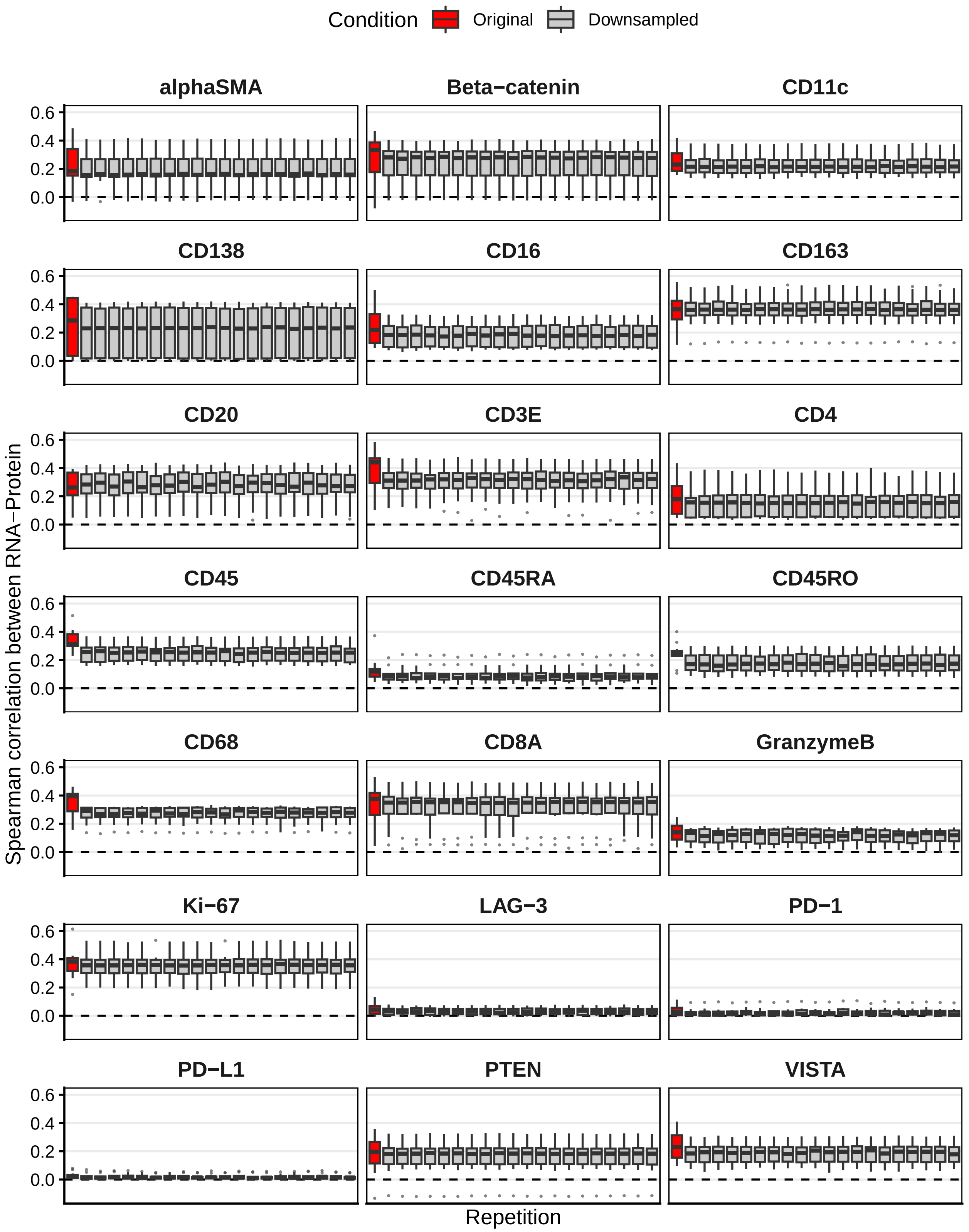


**Figure S5: Transcript down sampling stability.** Distribution of spearman correlation concordance between RNA and protein across all mCRPC samples, at original transcript depth (red) and across 20 downsampling simulations (gray).

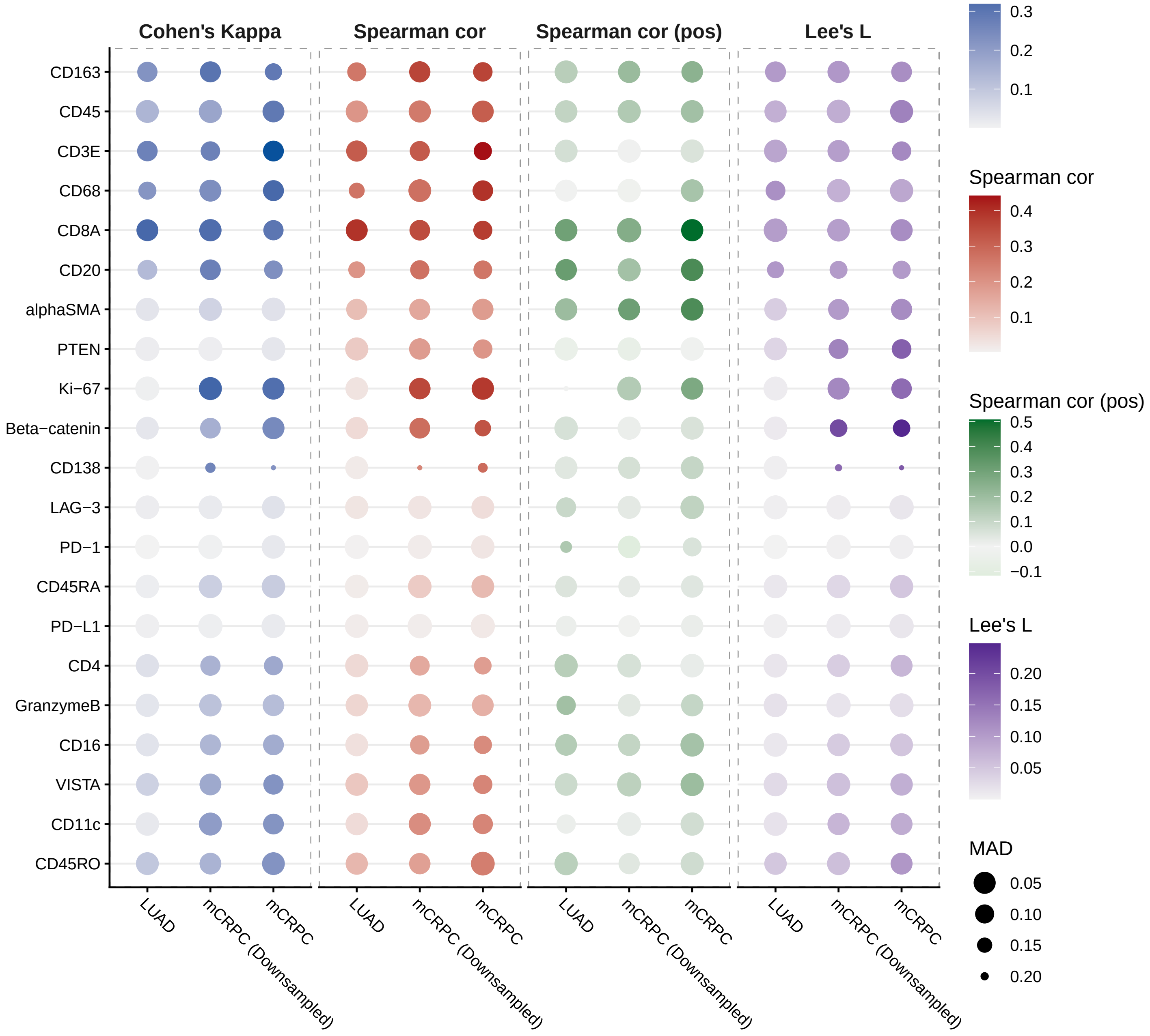


**Figure S6: Concordance metrics summary.** Dot plot summarizing RNA–protein concordance in each dataset. The evaluated metrics were detection agreement (Cohen's κ), Spearman's correlation across all cells, Spearman's correlation among positive cells, and Lee's L spatial concordance. Color indicates the median value of each concordance metric, and point size indicates its median absolute deviation (MAD).

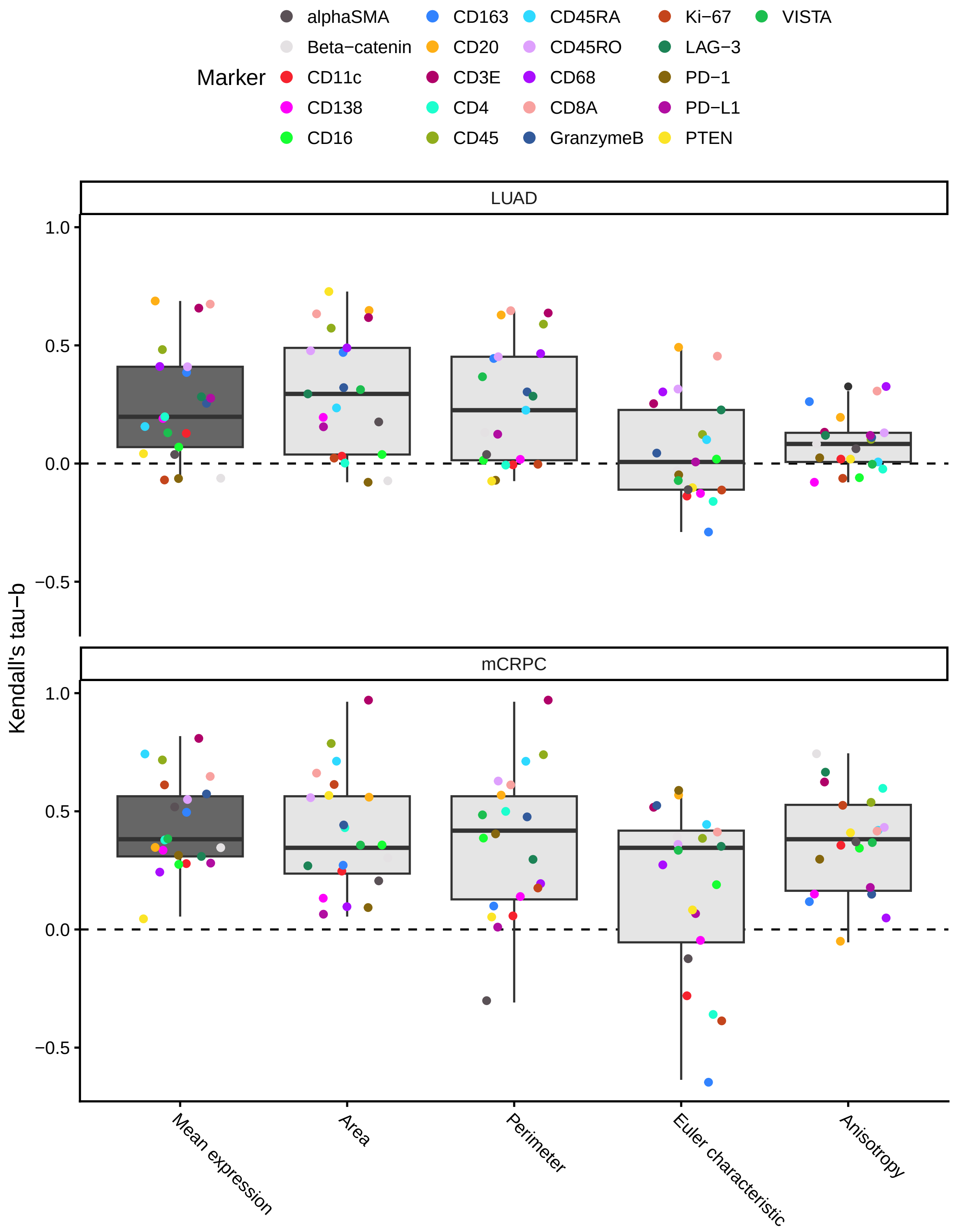


**Figure S7: Sample-ranking concordance.** Concordance across summary metrics. Boxplot of Kendall's tau-b coefficients quantifying rank concordance between modalities across samples for each metric: mean expression (pseudobulk), area, perimeter, Euler characteristic, and anisotropy (spatial). Each box shows the distribution of τ-b values across protein markers (colored individually) within that metric.
